# Injury-Free In Vivo Delivery and Engraftment into the Cornea Endothelium Using Extracellular Matrix Shrink-Wrapped Cells

**DOI:** 10.1101/2021.03.01.433476

**Authors:** Rachelle N. Palchesko, Yiqin Du, Moira L. Geary, Santiago Carrasquilla, Daniel J. Shiwarski, Irona Khandaker, James L. Funderburgh, Adam W. Feinberg

## Abstract

Cell injection is a common clinical approach for therapeutic delivery of cells into diseased and damaged tissues in order to achieve regeneration. However, cell retention, viability, and engraftment at the injection site has generally been poor, driving the need for improved approaches. Here, we developed a technique to shrink-wrap micropatterned islands of corneal endothelial cells in a basement membrane-like layer of extracellular matrix (ECM) that enables the cells to maintain their cell-cell junctions and cytoskeletal structure while in suspension. These μMonolayers exhibited the ability to rapidly engraft into intact, high-density corneal endothelial monolayers in both in vitro and in vivo model systems. Importantly, the engrafted μMonolayers increased local cell density, something that the clinical-standard single cells in suspension failed to do. These results show that shrink-wrapping cells in ECM dramatically improves engraftment and provides a potential alternative to cornea transplant when low endothelial cell density is the cause of corneal blindness.

**One Sentence Summary:** Shrink-wrapped patches of endothelial cells can rapidly attach and integrate into an intact cornea endothelium when injected into the anterior chamber, increasing cell density.

## Introduction

Organ and tissue transplants are the only option for patients with end-stage organ failure, and while effective, >1 million people are unable to benefit due to global donor shortages (*1*). Patients must also remain on immunosuppressants for life with major side-effects, experience a high rate of organ failure and rejection, and have no access to transplantation in many parts of the world (*2*). As an alternative, cell-based therapies have long been thought of as a potential therapeutic option for a range of diseases and injuries caused by tissue and organ failure such as myocardial infarction (*3*), diabetes (*4, 5*), corneal blindness (*6*), cystic fibrosis (*7, 8*). The goal is to deliver viable cells that integrate into the target tissue and replace damaged or dysfunctional cells in order to stop or reverse disease progression. Potential advantages compared to transplant include minimally invasive cell delivery without the need for extensive surgery, use of autologous cells to avoid immune rejection, and the ability to improve tissue and organ functional earlier in the disease process and thus entirely avoiding end-stage failure. Indeed, the past decade has seen advances in research and development of cell-based therapies based on autologous adult stem cells and induced pluripotent stem (iPS) cells (*9*–*11*). However, simple injection of cells into tissues has shown only limited clinical success in many applications due to low cell viability after injection as well as poor retention at the injection site and engraftment into the damaged tissue (*10, 12, 13*). Thus, there remains critical need for new technologies that can improve cell delivery, engraftment, and function.

The cornea serves as a clinically relevant tissue for development of new cell delivery approaches because at >50,000 procedures annually in the US, it is transplanted more than all other solid organs combined (*14*). Specifically, here we are focused on the corneal endothelium (CE), a single layer of cells that lines the posterior surface of the cornea and is responsible for maintaining proper corneal thickness and clarity through regulation of stroma hydration. Nearly 50% of all corneal transplants are due to failure of the CE, primarily due to loss of CE cells that are cell cycle arrested and cannot replicate to repair damage or injury (*15*–*18*). This subsequently leads to failure to properly pump fluid from the stroma to the aqueous humor once the cell density drops below ∼500 cells/mm^2^, resulting in corneal edema and clouding (*19, 20*). Current clinical treatment for CE failure is full thickness penetrating keratoplasty (PK) or partial-thickness transplants such as Descemet membrane endothelial keratoplasty (DMEK) and Descemet stripping automated endothelial keratoplasty (DSAEK) (*14, 21*). These lamellar techniques have shown improvement over PK, with evidence that immune rejection is reduced with less stroma and extracellular matrix (ECM) transplanted (*22, 23*). Further, the eye is considered to be immune privileged and eye drops are usually adequate rather than systemic immunosuppression. However, chronic rejection and limited donor supply in many parts of the world have motivated the development of new methods to inject CE cells into the anterior chamber to repopulate the endothelium and restore function (*6*). The problem these cell therapies have faced in the eye is the same as in other tissues and organs, effective delivery, and engraftment (*10, 12, 13*). In fact, most approaches require the existing CE to be removed through scrapping or cryogenic injury of the cornea in order to provide a place for the delivered cells to attach (*6*).

Here we report development of a new cell delivery method designed to enhance cell attachment and engraftment into tissues in vivo without requiring any induced damage to achieve integration. The challenge to delivering cells to the CE, and to epithelial and endothelial layers in general, is that these tissues are characterized by robust cell-cell injunctions and in general have evolved to act as barrier to keep things out. Thus, it has proved challenging to deliver single cells in suspension, which do not have a mechanism to attach and integrate. To address this, we hypothesized that small patches of CE with intact tight junctions and cytoskeletal structure may exhibit improved adhesion and integration into existing CE monolayers compared to single cells. Specifically, our goal was to create a method where we could deliver viable cells to intact CE monolayers without removal of any cells and achieve integration that would increase cell density. To do this we developed an approach to shrink-wrap micron-scale monolayers (μMonolayers) of CE cells within a engineered layer ECM using an adaptation of our previous reported surface-initiated assembly technique (*24, 25*). This technology enables the cells within the μMonolayers to maintain viability, tight-junctions and cytoskeletal structure throughout the release and injection process. Most importantly, the μMonolayers are able to integrate into existing CE monolayers and significantly increased cell density in both in vitro and in vivo assays. These results suggest that this technique could be used to increase cell density to treat corneal blindness without requiring the removal of the existing CE and enhance the engraftment of injected cells.

## Results

### Shrink-wrapped CE cells µMonolayers maintain cytoskeletal structure, tight junctions, and high viability

To engineer the CE cell μMonolayers, bovine or rabbit CE cells were seeded onto micropatterned 200 × 200 μm squares of ECM proteins (1:1 laminin and collagen IV) and ∼5 nm thick (**fig. S1**) that were fabricated via surface-initiated assembly on thermo-responsive poly(n-isopropylacrylamide) (PIPAAm) substrates (**Fig. 1**). The cells were cultured on the scaffolds for 24 hours to allow the cells to adhere to the ECM square, establish cytoskeletal structure and tight junctions, and form a confluent layer. Upon thermally triggered dissolution of the PIPAAm, the ECM square releases from the surface and effectively shrink-wraps around the CE cells forming the μMonolayer. Due to inherent pre-stress in the CE cells from being spread on the PIPAAm surface, once released the μMonolayers contract in size and are small enough to be injected through a small gauge needle.

**Fig. 1.**
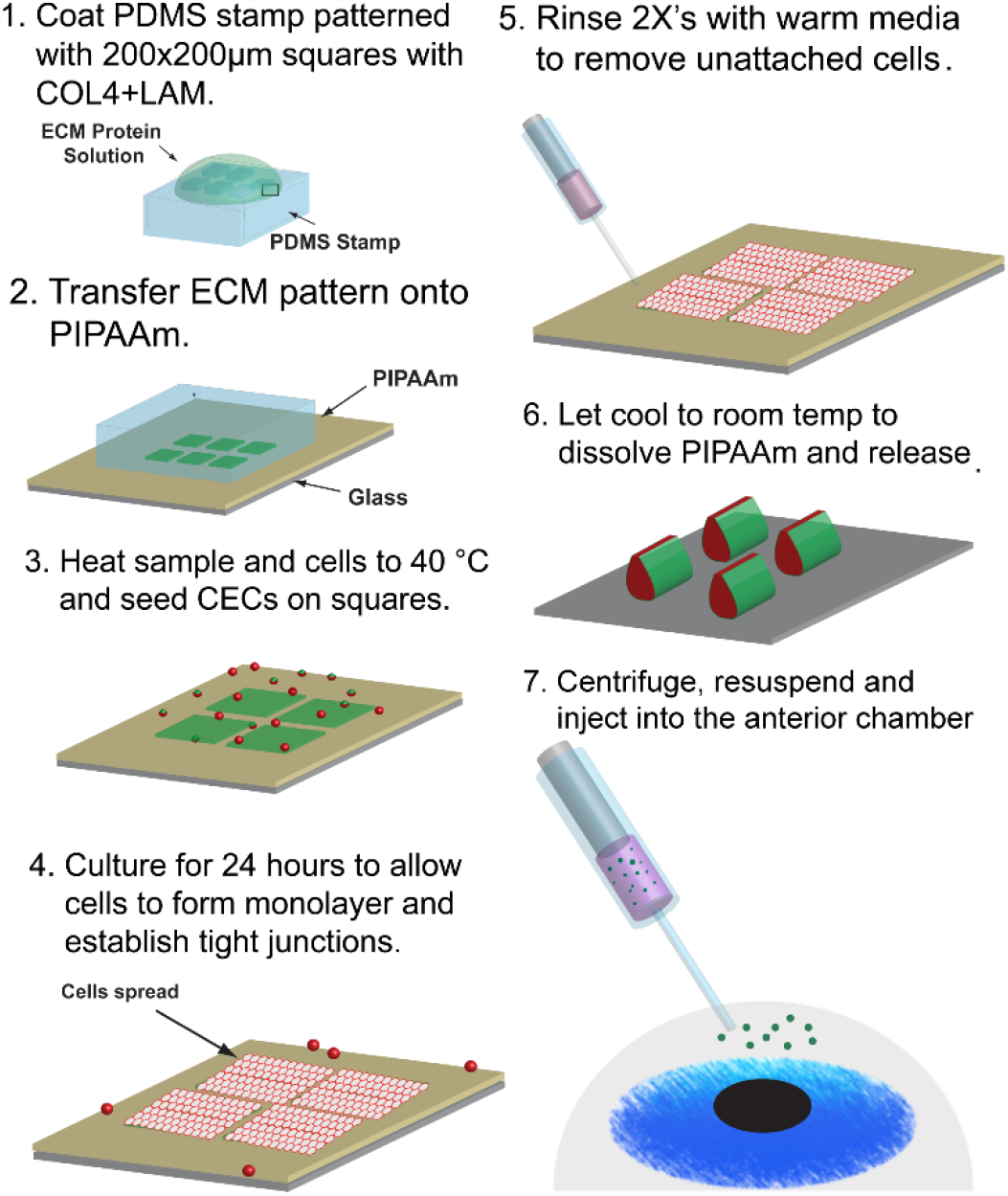
Schematic representation of the process for shrink-wrapping and injecting corneal endothelial cell μMonolayers. Steps 1&2: Surface initiated assembly techniques are used to engineer 200 μm square, 5 nm thick ECM scaffolds on the thermoresponsive polymer, PIPAAm. Steps 3&4: The samples and cells are then heated to 40 °C before seeding the cells on the squares and culturing for 24 hours. Steps 5&6: After 24 hours, samples are rinsed with warm media and cooled to room temperature to trigger the dissolution of the PIPAAm and shrink-wrapping/release of the µMonolayers of corneal endothelial cells before injection into the anterior chamber of the eye (Step 7).

To create the μMonolayers, we needed to ensure that the CE cells would adhere and spread on the ECM squares to form confluent patches and then properly shrink-wrap. As a control, bovine CE cells were seeded onto ECM squares micropatterned onto polydimethylsiloxane (PDMS) substrates because it is not temperature sensitive and previous studies have established cell growth on this surface (*25*–*29*). After 24 hours, the CE cells on the ECM squares on PDMS were adhered and spread into a monolayer as expected (**Fig. 2A**). Next, we repeated this by seeding CE cells on ECM squares patterned on PIPAAm, and the cells exhibited a similar morphology when viewed under phase microscopy (**Fig. 2B**). Upon dissolution of the PIPAAm and thermal release, the cells remained interconnected and were successfully shrink-wrapped within the ECM squares into µMonolayers (**Fig. 2C**). Time-lapse images show that once the media reached room temperature and the PIPAAm dissolved (0 sec), the shrink-wrapping process occurred quickly in <100 seconds (**Fig. 2D, video S1**).

**Fig. 2.**
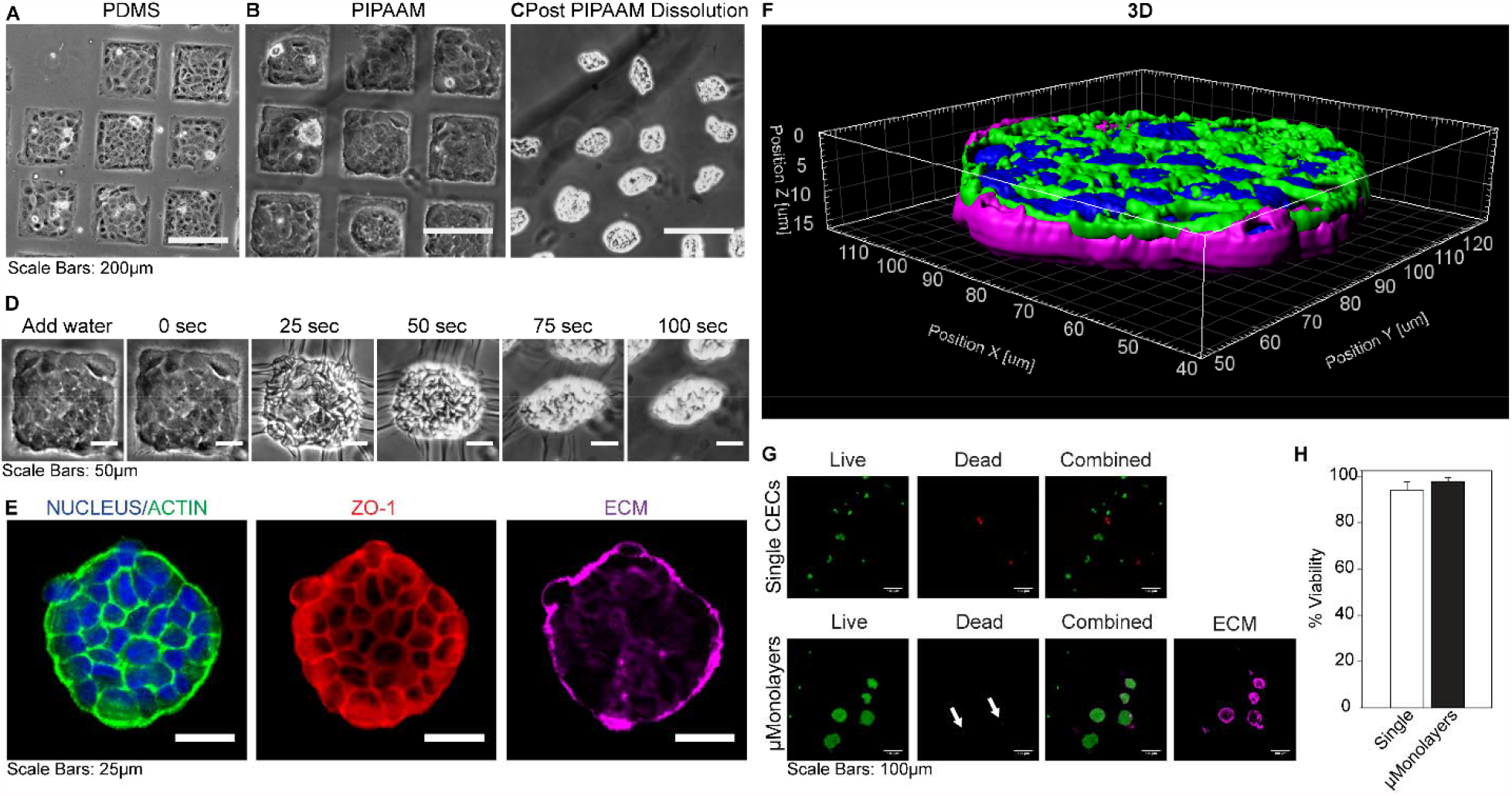
CE cells form μMonolayers on ECM squares and maintain their structure and viability through shrink-wrapping and injection. **(A)** CE cells form monolayers on ECM squares microcontact printed onto PDMS (used as a control) and **(B)** on the thermoresponsive polymer PIPAAm. **(C)** Once the PIPAAm is dissolved the CE cell μMonolayers contract and are shrink-wrapped in the ECM squares. **(D)** The release and shrink-wrapping of μMonolayers occurs quickly, in <100 seconds once the water + sample cools to room temperature. **(E)** Confocal microscopy images show that after injection, the CE cells maintain both their cytoskeletal structure (F-actin, green), tight junctions (ZO-1, red) and adherence to the ECM scaffold (LAM+COL4, purple). **(F)** A 3D projection of a shrink-wrapped CE cell μMonolayer 30 minutes after injection onto a glass surface illustrating how it begins to relax and return to its original shape. **(G)** Representative live/dead images of control single CE cells and shrink-wrapped CE cells show that both types of cells are viable with very few dead cells present. **(H)** Live/dead data showed no significant difference in viability between single cells (93 ± 4 %) and shrink-wrapped cells (97 ± 2 %) following injection through a 28G needle (n=3; mean ± stdev.; N.S. by Student’s t-test Single vs. μMonolayers).

After release, the shrink-wrapped µMonolayers were collected, centrifuged, injected through a 28G needle onto a glass coverslip and allowed to settle for 30 min before fixing and staining to investigate the morphology, structure, and viability of the CE cells. The shrink-wrapped CE cells exhibited continuous ZO-1 at the borders and a cortical F-actin structure indicating that the cells maintained their tight-junctions and cytoskeletal structure throughout the release and injection process (**Fig. 2E**). Immediately post-release, the shrink-wrapped µMonolayers contracted tightly into small clusters (**Fig. 2C**), however, approximately 30 minutes after release and injection the μMonolayers relaxed and returned to a disc-like morphology as they settled onto the surface (**Fig. 2F**). This establishes that the shrink-wrapping process and subsequent injection through a small gauge needle does not disrupt cell-cell adhesions or cause damage to the CE cells in the μMonolayer. High cell viability in the µMonolayers after injection was confirmed using a Live/Dead cytotoxicity assay and compared to enzymatically released single cells (**Fig. 2G**). Confocal microscopy images revealed that the only dead cells were those that were not integrated into the shrink-wrapped µMonolayers, with cells in the μMonolayers showing very high viability (**Fig. 2G**, shown by the arrows). Quantitative image analysis showed that single CE cells in suspension had 93 ± 4% viability and the CE cells within the shrink-wrapped μMonolayers had 97 ± 2% viability, though this difference was not statistically significant (**Fig. 2H)**.

### Shrink-wrapped µMonolayers rapidly adhere and spread to form a CE monolayer on collagen gels in vitro

To assess the potential of the shrink-wrapped µMonolayers for cell injection therapy, we first performed an in vitro assay using a compressed collagen type I gel as a model of a denuded corneal stroma. Shrink-wrapped µMonolayers and single CE cells in suspension (as the control) were seeded onto compressed collagen type I gels by injecting through a 30-gauge needle. Samples were fixed and stained post-injection at 6 hours to observe initial attachment and adhesion and at 24 hours to observe and spreading and outgrowth. At 6 hours post-injection, single CE cells were mostly rounded with very little spreading observed (**Fig. 3A**) and the F-actin staining showed a lack of filamentous cytoskeleton structure, additionally there was no ZO-1 observed. In contrast, the CE cells from the shrink-wrapped µMonolayers maintained their cytoskeletal structure and tight-junctions, as evidenced by the F-actin filaments and continuous ZO-1 expression at the cell borders (**Fig. 3B**). Examining the samples in 3D confirmed that single CE cells were rounded and had few contacts between cells (**Fig. 3A**) whereas the shrink-wrapped µMonolayers had reoriented with cells directly attached to the collagen and the ECM scaffolds now present within the center of the monolayer (**Fig. 3B**). After 24 hours, the single CE cells covered most of the collagen substrate and had a more defined cytoskeletal structure (**Fig. 3C**) but with many F-actin stress fibers across the cell bodies rather than being primarily cortical. Additionally, the single CE cells exhibited very low ZO-1 staining, expressed discontinuously at the cell borders (**Fig. 3C**). In contrast, the shrink-wrapped µMonolayers had continuous ZO-1 at all cell borders and abundant cortical F-actin (**Fig. 3D**), which closely resembled the structure of in vivo CE cells (*30*). The remnants of the ECM squares that had shrink-wrapped the are visible as indicated by the arrows in **Fig. 3D**. These results show that CE cells in shrink-wrapped μMonolayers more rapidly repopulate a collagen substrate than CE cells in suspension over the first 24 hours of culture.

**Fig. 3.**
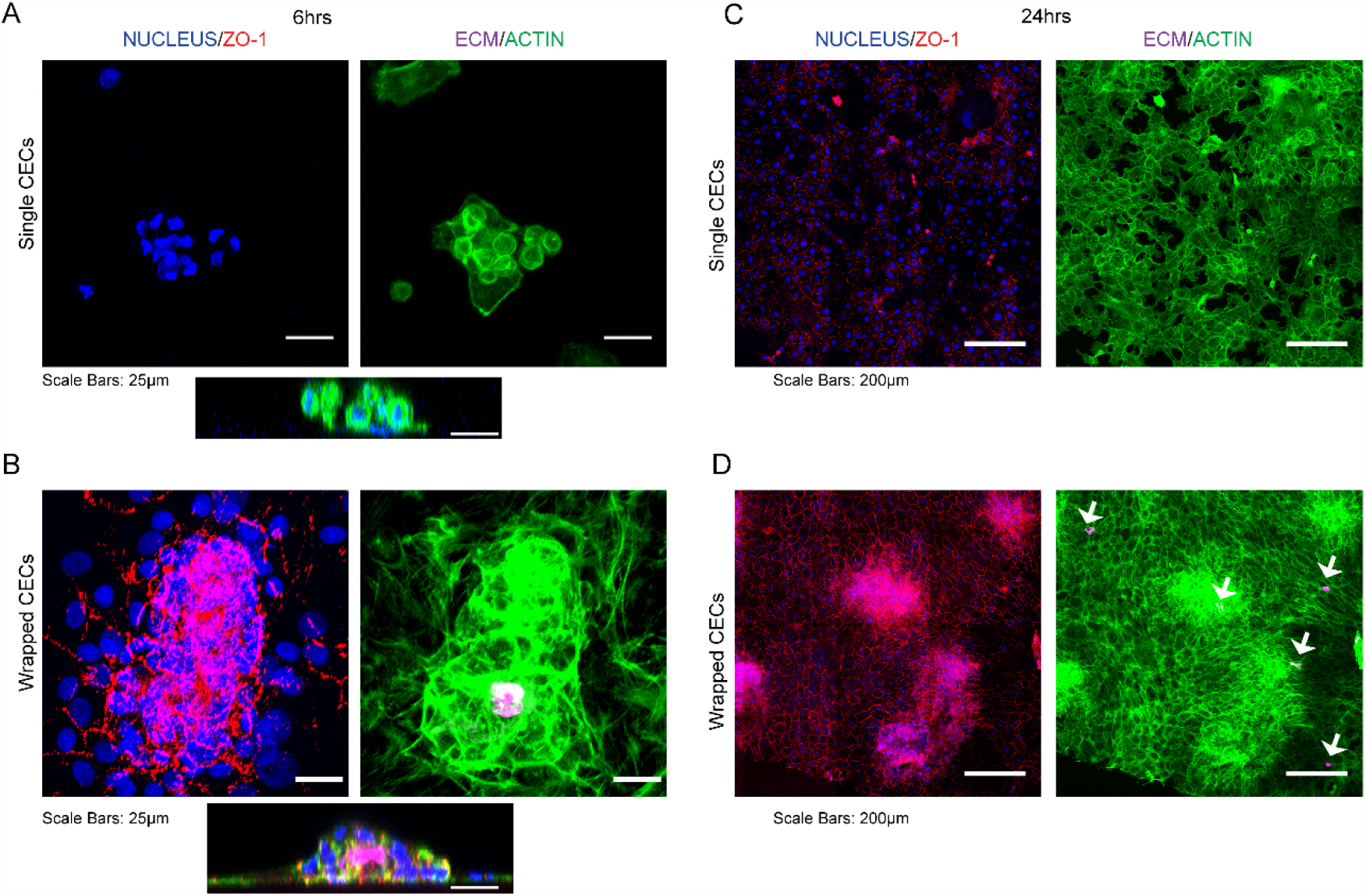
Shrink-wrapped CE cell μMonolayers maintain ZO-1 expression F-actin cytoskeleton as they grow out of the ECM scaffolds to form a monolayer on a collagen type I stromal mimic. **(A)** Six hours after reseeding onto a collagen type I gel, the single CE cells have no established F-actin cytoskeleton or ZO-1 expression. In contrast, the CE cells in the shrink-wrapped μMonolayers have maintained their ZO-1 expression and F-actin cytoskeleton, while growing out of the ECM scaffolds. The cells at the periphery of the shrink-wrapped CE cells are also expressing ZO-1. **(B)** The 3D views of the cells at 6 hours post-seeding show the differences between the single CE cells and shrink-wrapped CE cells. Images on the left are cross-sectional projections of the 3D views. **(C)** At 24 hours, single CE cells have begun to spread and cover almost the entire scaffold. **(D)** At 24 hours, the CE cells have already grown out of the ECM scaffolds and formed an almost complete monolayer. For A-D: Nucleus = blue, ZO-1 = red, ECM(COL4) = magenta, F-actin =green.

### Shrink-wrapped µMonolayers show enhanced engraftment into existing CE monolayers and increase CE density in vitro

There are currently clinical trials in Japan using human induced pluripotent stem cell (iPSC) derived CE cells to restore the endothelium, but it requires removal of the existing CE cells in order to make space to deliver the new cells and allow them to attach(*6, 31*). This is because cells seeded onto an existing endothelial or epithelial layer typically show very low attachment and engraftment. Rather than damaging a tissue in order to repair it, we hypothesized that the shrink-wrapped µMonolayers would be able to engraftment into existing CE monolayers through enhanced cell-cell and cell-ECM binding. To test this, we seeded shrink-wrapped CE cell µMonolayers in vitro onto a low-density CE monolayer to mimic that observed in patients that need a cornea transplant. The CE cells were labeled with CellTracker green and µMonolayers or as a single cell suspension control were injected through a 30G needle to seed them. CE monolayers that were not seeded with any cells served as negative controls. The seeded CE single cells and µMonolayers were allowed to settle onto the samples for 3 hours before rinsing and adding fresh media to mimic the procedure used for clinical CE cell injection in animal models and in human patients, where they remain face down for 3 hours post-injection (*6, 32, 33*). Samples were then cultured for 3, 7 and 14 days and analyzed for engraftment (**Fig. 4A**). At each time point, very few single CE cells were integrated into the CE monolayers, and large areas of the samples had to be imaged to find even a few labeled cells. In contrast, the shrink-wrapped µMonolayers were well integrated at day 3 and appeared more densely packed compared to the original monolayer. At day 7, the shrink-wrapped CE cells had completely integrated into the existing CE monolayer, still appearing more densely packed but also more spread out (**Fig. 4A**). By day 14, the µMonolayers appeared to be well engrafted and the density of the newly integrated CE cells had equilibrated with the CE cells in the existing monolayer, achieving an increased overall cell density.

**Fig. 4.**
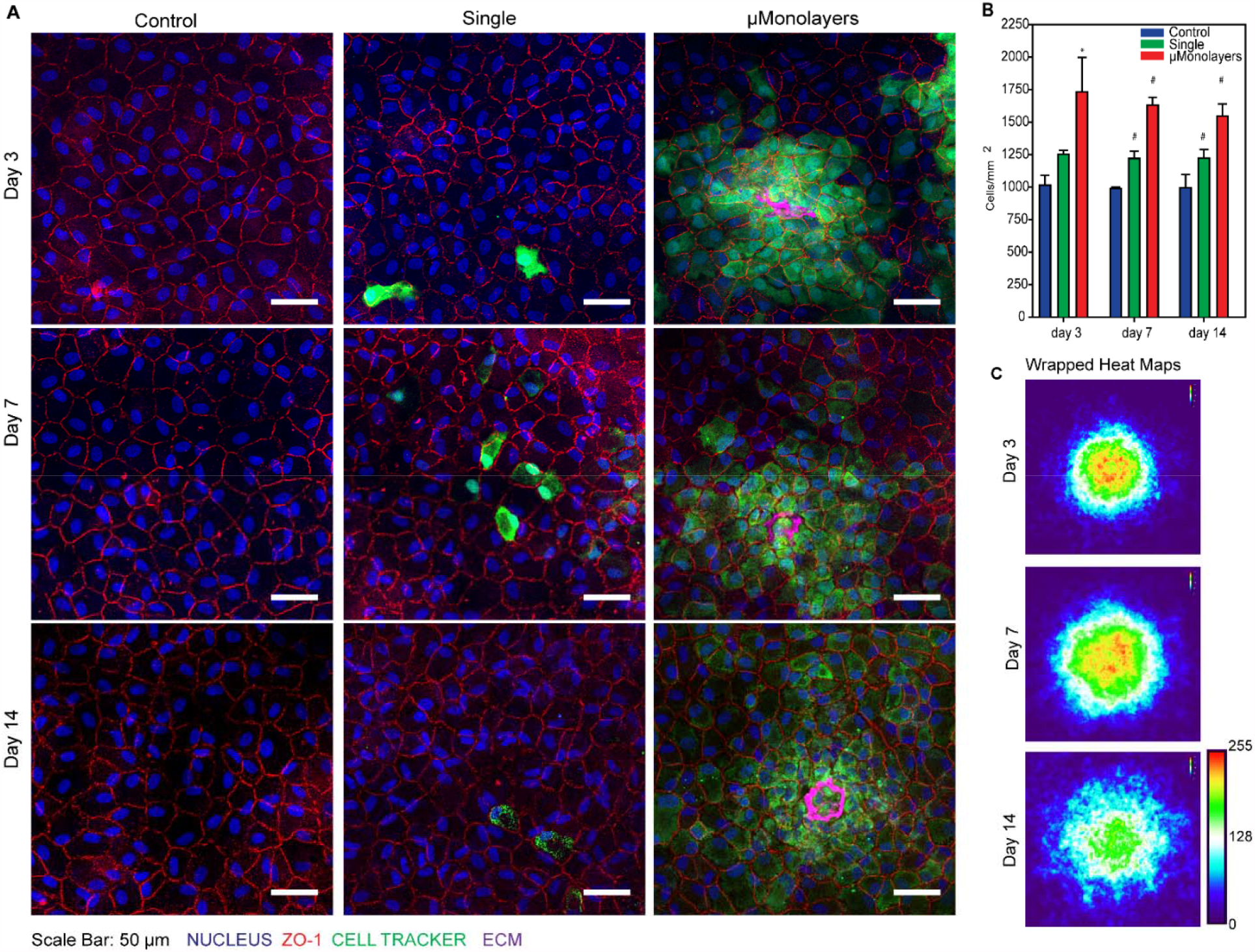
Injected shrink-wrapped μMonolayers integrate into existing monolayers of CE cells and significantly increase the density compared to single CE cells. **(A)** Cell Tracker labeled single cells and shrink-wrapped cells were visible at all time points however, significantly more shrink-wrapped cells were present at all time points and the ECM scaffolds were still visible 14 days after injection. Scale bars = 50 μm. **(B)** The cell density of the monolayers was calculated and compared at days 3, 7 and 14. Data is represented as mean ± std dev. Day 3: Control = 1016 ± 75 cells/mm^2^; Single = 1253 ± 31 cells/mm^2^; μMonolayer = 1731 ± 267 cells/mm^2^. Day 7: Control = 989 ± 11 cells/mm^2^; Single = 1220 ± 56 cells/mm^2^; μMonolayer = 1631 ± 58 cells/mm^2^. Day 14: Control = 994 ± 104 cells/mm^2^; Single = 1224 ± 66 cells/mm^2^; μMonolayer = 1545 ± 95 cells/mm^2^.The data was compared using a one-way ANOVA on ranks with Tukey’s test (day 3) or one-way ANOVA (day 7 and 14) with Tukey’s Test is SigmaPlot. * = statistically significantly different from control, # = statistically significantly different from all other samples. Day 3: n=4 for all samples; Day 7: control n=3, single n=4, wrapped n=4; Day 14: control n=4, single n=3, wrapped n=4. **(C)** Heat maps of Cell Tracker positive pixels show that the cells in the shrink-wrapped μMonolayers initially integrate into a tight cluster and then the density equilibrates as the cells spread out slightly (Day 3 n= 33, Day 7 n= 37, Day 14 n= 40, scale bar is arbitrary units).

To determine how well the shrink-wrapped CE cell µMonolayers engrafted and increased the cell density of the low-density CE monolayers, the density was analyzed at each time point. Overall, the single CE cells increased the monolayer density ∼20% while the shrink-wrapped CE cell µMonolayers increased the monolayer density ∼50%, even though the same total number of cells were seeded for each condition (**Fig. 4B**). At day 3, the shrink-wrapped µMonolayers significantly increased cell density compared to controls and at days 7 and 14 the shrink-wrapped µMonolayers significantly increased cell density compared to both controls and samples seeded with single CE cells. These results establish that the shrink-wrapped µMonolayers adhere, engraft, and then spread out into an existing CE monolayer in a manner that single CE cells cannot. Further, we generated heat maps using images of the CellTracker green labeled CE cells in the shrink-wrapped µMonolayers to confirm that the cells were spreading out over time (**Fig. 4C**). The results confirm that at day 3 the CE cells are contained in a small area and that over time they spread out into the surrounding monolayer and equilibrate in density. This makes sense, because even though the images of fixed CE monolayers makes it look like the cells are stationary, time-lapse images routinely show that cells are constantly moving within epithelial layers (*34, 35*), which should facilitate the spreading out of the shrink-wrapped CE cells. This equilibration process also explains the perceived decrease in cell density from day 3 to 14, where the cell density overall isn’t decreasing, it is just becoming more homogenous across the CE monolayer. This was confirmed by running a one-way ANOVA within each sample type comparing the densities over time, which showed that the time point had no statistically significant effect.

### Shrink-wrapped µMonolayers adhere and begin to integrate into the CE within 3 hours

Clinical injection of single CE cells requires that patients lie face down for 3 hours post injection to allow for cell attachment to the CE on the posterior of a denuded cornea (*6, 31, 32*). Thus, a major question for the shrink-wrapped μMonolayers is how they achieve improved engraftment and how long does it take compared to single cells. To address this, we performed live confocal imaging of engraftment in vitro by labeling the shrink-wrapped μMonolayers with CellTracker™ Green and the cells in the low-density monolayer with CellTracker™ Orange. Note that we performed this study in vitro because we could take time lapse confocal images, which would not be feasible in an animal model. By collecting a Z-stack every hour for 48 hours, we were able to clearly observe the dynamics of the integration process from both top-down and side views (**videos S2 and S3**). At 3 hours post-injection, the shrink-wrapped µMonolayers had begun to attach and flatten on top of the low-density monolayer (**Fig. 5A**). Over time the shrink-wrapped µMonolayers continued to flatten and move around as the cells moved into the underlying low-density monolayer. By 43 hours, the shrink-wrapped µMonolayer was almost completely integrated into the CE monolayer and appeared to be in the same imaging plane with the low-density monolayer and not sitting on top of it. The ECM square used in the shrink-wrapping process remained centrally located underneath the cell bodies of the shrink-wrapped cells, which is consistent with results from the fixed time point experiments (**Fig. 4A**).

**Fig. 5.**
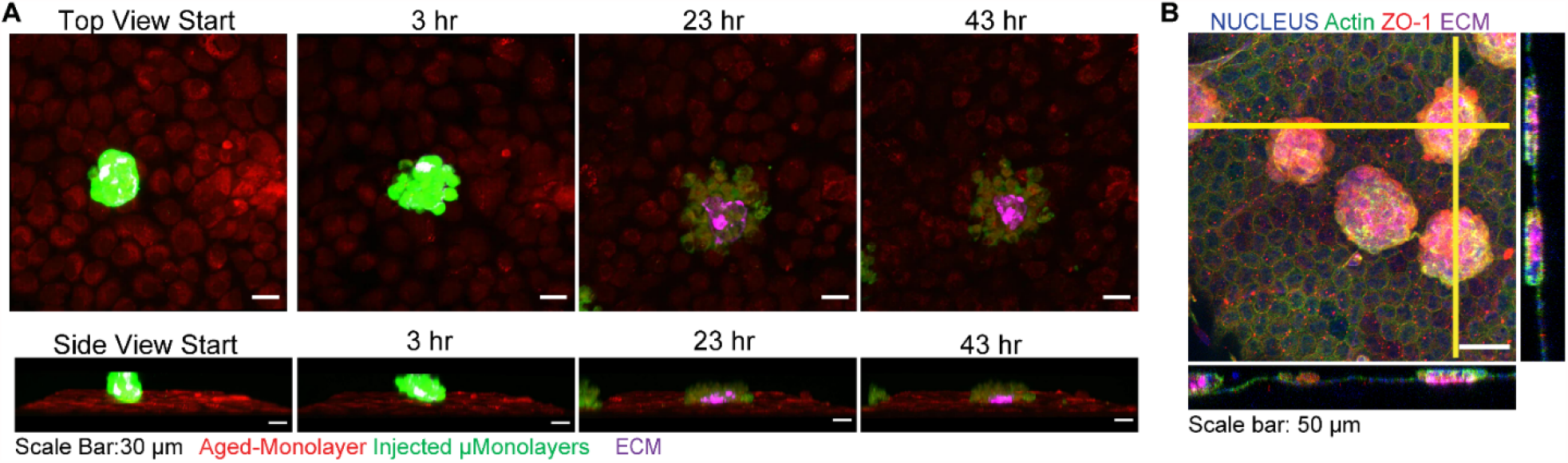
Shrink-wrapped μMonolayers begin to integrate into “aged” CE monolayers and ex vivo corneas within 3 hours. **(A)** Time-lapse images from live confocal imaging of the integration of shrink-wrapped bovine μMonolayers (Cell Tracker green) into an engineered “aged” bovine CE monolayer (Cell Tracker Orange). At 3 hours, the μMonolayers have attached and begun to integrate and by 43 hours the cells are almost completely integrated into the monolayer. **(B)** Confocal images show that the shrink-wrapped μMonolayers had begun to integrate into the ex vivo rabbit CE and the ECM scaffold is observed to be between μMonolayers and the existing rabbit CE. The yellow vertical and horizontal lines indicate the places at which the orthogonal views were obtained.

Although the time-lapse imaging results suggested that 3 hours is sufficient for the shrink-wrapped µMonolayers to attach to the CE, potential differences between in vitro and in vivo conditions caused us to assess attachment to the native cornea. To do this, we switched to rabbit CE cells and injected shrink-wrapped µMonolayers ex vivo into the anterior chamber of enucleated rabbit eyes. The eyes were then incubated with the cornea facing down for 3 hours to allow attachment of the shrink-wrapped µMonolayers before fixation and staining of the whole globe (**fig. S2**). Confocal imaging of the corneas showed numerous shrink-wrapped µMonolayers attached over the entirety of the posterior surface (**Fig. 5B**). At this 3-hour time point, the shrink-wrapped µMonolayers were still disc-like in shape and oriented with the ECM layer facing the CE on the posterior surface of the cornea. This is consistent with the observations from the time-lapse experiments. These experiments provided confidence that 3 hours was sufficient for gravitational settlement and attachment of the shrink-wrapped µMonolayers for subsequent in vivo experiments.

### Shrink-wrapped µMonolayers show robust engraft into the CE in vivo

Having demonstrated enhanced engraftment of the shrink-wrapped µMonolayers in vitro and ex vivo, we moved next to an in vivo rabbit model. First, we assessed the basic feasibility of injecting the shrink-wrapped µMonolayers in vivo and achieving engraftment. To do this, shrink-wrapped µMonolayers (n=3) or single cells (n=2) were labeled with DiO and 100,000 cells in 50 μL of DMEM/F12 were injected into the anterior chamber of one eye for each rabbit. The rabbits were laid on their sides with the injected eye facing down for 3 hours to allow for cell attachment and integration and then followed daily for 1 week before sacrifice and enucleation. At 1 week, the injected eyes on all 5 rabbits remained clear with no visible outward signs of irritation or swelling and appeared to be same as the contralateral control eye in each animal (**Fig. 6A,B, fig. S3**.). Additional examination by Confoscan indicated no abnormalities in the corneal endothelium (**fig. S4**). These results establish that (i) the injection process to deliver shrink-wrapped µMonolayers to the anterior chamber does not damage the cornea, and (ii) that there is no immune response in terms of cell infiltration that would cloud the cornea due to the allogeneic rabbit CE cells.

**Fig. 6.**
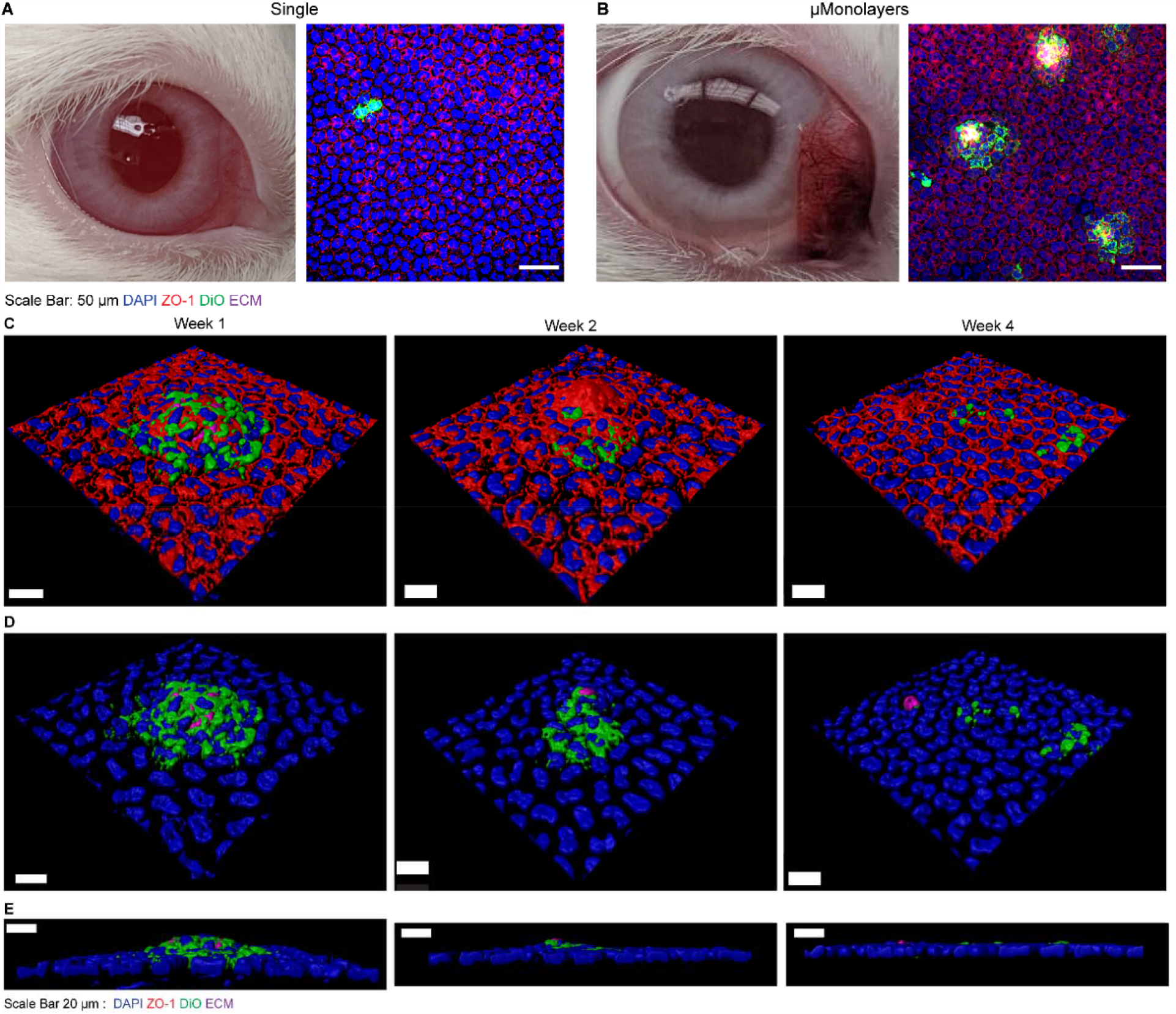
Shrink-wrapped μMonolayers integrate into the existing healthy rabbit CE. **(A)** Rabbit corneas injected with single cells remained clear at 1-week post injection. However, very few DiO labeled single cells are observed integrated into the rabbit CE. **(B)** The rabbit cornea injected with μMonolayers also remained clear 1-week post injection and numerous clusters of μMonolayers were observed in each cornea with the continuous ZO-1 at the borders between DiO labeled cells and native rabbit CE cells. **(C)** Graph showing the cell density in the areas of the integrated shrink-wrapped μMonolayers (green bar) compared to native areas within the same image with no DiO labeled cells (blue bar). Data is represented as mean ± standard deviation and was compared using a student T-test. It was found that the density in the areas with shrink-wrapped μMonolayers was significantly higher than the native CE cell density (* p<0.05). **(D)** Confocal microscopy images showing the integrated DiO labeled shrink-wrapped μMonolayers at 1, 2- and 4-weeks post-injection. Nuclei = blue, DiO labeled cells (green), ZO-1 (red), ECM nanoscaffold (purple). **(E)** The same images from panel D with the ZO-1 removed to highlight the nuclei of both the healthy rabbit endothelium and injected cells (blue), DiO labeled injected rabbit cells (green) and ECM nanoscaffold (purple) from the shrink-wrapping process at 1, 2- and 4-weeks post-injection. **(F)** The orthogonal views of the confocal images shown in panel E showing the integration of the shrink-wrapped μMonolayers into the healthy rabbit endothelium over the 4-week period post-injection. Scale bars in D-F are 20 μm.

After enucleation, the eyes were fixed, and the corneas were stained as wholemounts for the tight junctions via ZO-1 and the nuclei to observe integration into the healthy rabbit CE. Confocal microscopy imaging showed that very few DiO labeled cells were present in the single cell injected eyes of both rabbits (**Fig. 6A and fig. S6A**) with only a few labeled cells being found across the entire cornea. In contrast, numerous clusters of shrink-wrapped µMonolayers integrated throughout the corneas of all 3 rabbits injected with the μMonolayers (**Fig. 6B and fig. S6B**). Higher magnification imaging showed that the cells of the shrink-wrapped µMonolayers had integrated with the healthy rabbit CE with ZO-1 present continuously at all cell borders between the DiO labeled cells and the native rabbit CE cells (**Fig. 6C**). Additionally, the ECM scaffolds were still visible, providing further evidence that the shrink-wrapped µMonolayers had integrated and become a part of the rabbit CE in vivo (**Fig.6C-E**).

To determine if the shrink-wrapped μMonolayers remained stable over time and to see if the high-density areas began to spread across the cornea, we next looked at the integration of the shrink-wrapped μMonolayers at 2 weeks (n=2) and 4 weeks (n=3). Comparable to week 1, all eyes at weeks 2 and 4 had no visible signs of irritation or swelling (**fig. S3** and Confoscan indicated no abnormalities in the corneal endothelium (**fig. S4**). At each time point labeled CE cells and ECM squares were detected, indicating that the cells remained viable, and stably integrated. Further, the presence of the ZO-1 between the native and shrink-wrapped cells indicated that a continuous monolayer was established (**Fig. 6C&D**). Interestingly, when the ZO-1 and F-actin channels are removed from the images, at week 1, the ECM appears to be at the center of the cell nuclei (**Fig. 7A**) but is also underneath of the cell bodies as the cells were not on top of the native CE cells and the continuous ZO-1 indicated the cells were integrated with the native cells. By weeks 2 and 4, the density of the nuclei tightly surrounding the ECM begins to decrease as the ECM also becomes smaller (**Fig. 6C, D & Fig. 7A**). The orthogonal views in **Fig. 6E** further confirms that the ECM begins to become smaller, and eventually the ECM and shrink-wrapped μMonolayer cells are flush with the surrounding native CE.

**Fig. 7.**
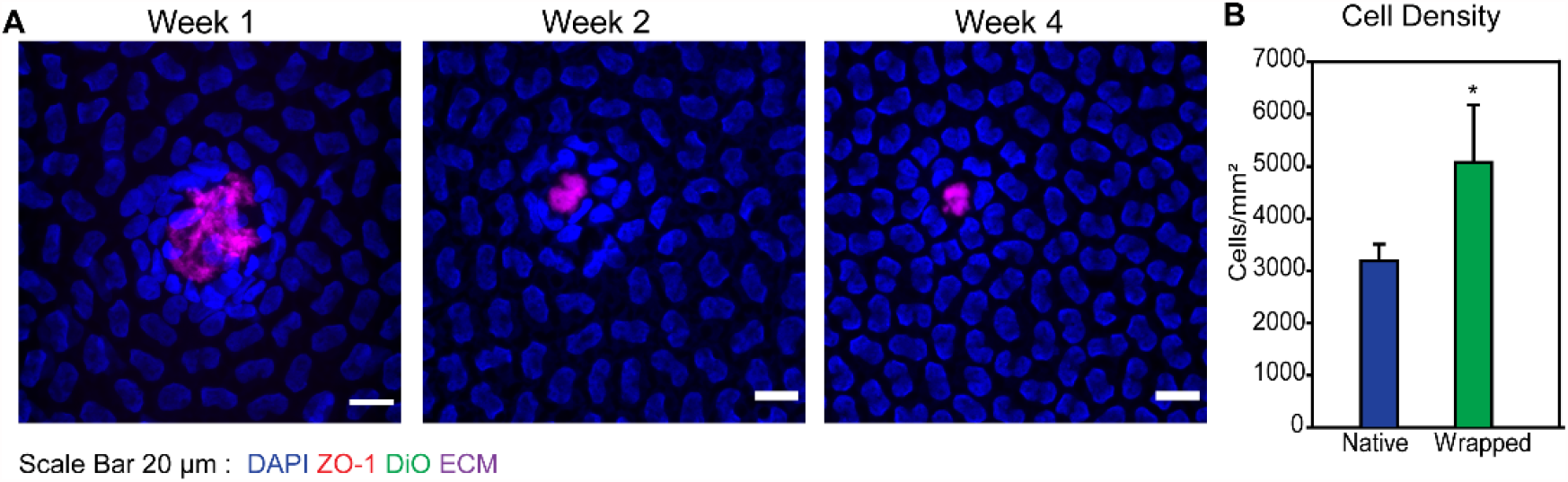
Shrink-wrapped μMonolayers exhibit stable integration into the existing healthy rabbit CE. **(A)** Confocal microscopy images showing the integrated shrink-wrapped μMonolayers at 1, 2- and 4-weeks post-injection. The images show that the ECM scaffold is under the cell bodies post integration and that the cell density around the ECM at 1 week post injection is higher and then begins to dissipate into the outer areas by week 4 post injection. **(B)** Graph showing the cell density in the areas of the integrated shrink-wrapped μMonolayers (green bar) compared to native areas within the same image with no DiO labeled cells (blue bar). Data is represented as mean ± standard deviation and was compared using a student T-test. It was found that the density in the areas with shrink-wrapped μMonolayers was significantly higher than the native CE cell density (* p<0.05).

Finally, we quantified the ability of the CE cells from the injected μMonolayers to stable integrate over time and increase the local cell density where they engrafted. At 1 week, cell density at locations with engrafted μMonolayers was compared to control areas within the same cornea (example areas shown in **fig. S5**). Results showed that the areas where the shrink-wrapped μMonolayers were attached an integrated had a significantly greater cell density compared to the surrounding areas of the native endothelium without shrink-wrapped μMonolayers (**Fig. 7A&B**). At 2- and 4-week time points, larger image areas were used as there is a possibility that the DiO fades with time due to (i) degradation of the dye and (ii) cell division that dilutes the DiO stain between two daughter cells. Therefore, we cannot be certain that all injected cells are still detectably fluorescing green. At 2 weeks, there was no significant difference in the density of the areas with DiO labeled cells compared to areas with no labeled cells (**fig. S7A**) and one of the 4-week eyes had a statistically significant difference in the cell density of areas with labeled cells (**fig. S7B**). These results, combined with the orthogonal views from the confocal imaging (**Fig. 6C-E**), indicate that by 2 weeks the CE density has begun to equilibrate across the entire native endothelium and there are no longer the higher density patches where the μMonolayers initially engrafted. These results are consistent with the engraftment and integration of CE cells from the shrink-wrapped μMonolayer experiments performed in vitro (**Fig. 4**). This equilibration of the CE density across the monolayer over times is in fact the behavior that we want to see, as it shows that the CE cells delivered to the cornea appear to behave as normal CE cells. Importantly, all of the in vivo results combined show that shrink-wrapped μMonolayers are able to stably engraft, integrate into the rabbit corneal endothelium, and remain viable long-term.

## Discussion

Based on our results and other research in the field, we put forward that delivery of cells that retain their phenotypic cytoskeletal structure enhances engraftment through increased cell-cell and cell-ECM adhesions. Indeed, the clinical translation of cell injection therapy into many tissues using single cells suspended in solution has faced a number challenges including limited or poor cell viability, retention, and integration at the injection site (*10, 12, 13*). In the literature, methods to improve cell injection therapy have generally focused on increasing either the retention or adhesion of the cells at the injection location using hydrogels, small molecules in the injection media (such as the ROCK inhibitor Y-27632), pre-conditioning of cells on ECM proteins, or genetic modification (*6, 33, 36*–*38*). While these methods have shown some improvements, they still fail to address the fact that single cells in solution have a rounded morphology that lacks the phenotypic cytoskeletal structure and cell-cell and cell-ECM interactions of these cells in their normal tissue microenvironment. Thus, at the time of injection single cells cannot immediately interact with their environment, and must actively reestablish their cytoskeleton, cadherins and other receptors to actively bind neighboring cells, and integrin to bind the ECM. In contrast, cells in shrink-wrapped μMonolayers when injected immediately start to interact with the surrounding cells and ECM, even being able to engraft into the challenging scenario of a confluent epithelial layer.

For example, previous work by the Kinoshita group on the culture and injection of CE cells has highlighted the significance of the actin cytoskeleton on the adhesion of CE cells in vivo. Their research has shown that enzymatic dissociation of cells induces the phosphorylation of myosin light chain (MLC) through the Rho/ROCK pathway, which induces actin contraction and this activation of MLC negatively regulates cell adhesion (*32*). To overcome this, they inject the ROCK inhibitor Y-27632 with the CE cells, which enhances cell adhesion by blocking the actin contraction, therefore increasing the interactions between the cytoskeleton and focal adhesion complexes and integrins (*33*). Cytoskeletal structure and tight-junctions are necessary for CE cell function, therefore, single cells may not be able to adhere to the existing monolayer and repair tissue level function of the damaged endothelial layer. We therefore hypothesized that (i) injected single cells have poor viability and attachment to intact tissues due to their lack of cell-cell junctions, cell-ECM interactions, and cytoskeletal structure and (ii) that monolayers of cells with dimensions small enough to be injected through a small gauge needle, would integrate into existing CE monolayers in higher numbers compared to single cells.

Our technology to shrink-wrap cells into µMonolayers using a thin layer of basement membrane-like ECM uniquely maintains high cell viability, cell-cell junctions, cell-ECM binding, and cytoskeletal structure post-injection. As shown, using the corneal endothelium as a model system these μMonolayers engraft into tissues within a few hours and established tight junctions and formed an organized F-actin cytoskeletal structure. When compared to enzymatically released single cells, injected shrink-wrapped µMonolayers engrafted at significantly higher numbers and increased CE monolayer cell density in vitro. Importantly, shrink-wrapped µMonolayers showed a high rate of engraftment into in vitro engineered low density monolayers, ex vivo corneas, and in vivo corneas. This was achieved without the need to disrupt cell-cell tight junctions or removal of the existing cells prior to seeding, which is the standard clinical practice for those patients with remaining CE cells (*6*). The in vivo rabbit studies were performed utilizing healthy rabbit eyes and results showed high numbers of shrink-wrapped µMonolayers integrated into the healthy CE and remained integrated over 4 weeks. This is extremely promising as cells within a young healthy rabbit endothelium are contact-inhibited, have tight-junctions, and are at an extremely high density. Therefore, if the shrink-wrapped µMonolayers can integrate within such a tissue, integration into damaged or diseased CEs with a much lower cell density would occur at much higher rates. This is all further evidence that injection of μMonolayers could be used for patients who are experiencing declines in cell density and visual acuity to boost their CE cell density before it reaches the lower limit where corneal blindness results, thus increasing the lifespan of their existing cornea and eliminating the need for a future transplant.

While these results are extremely promising, the in vivo study evaluated CE cell integration into a healthy, high-density endothelium that likely represents the most challenging engraftment environment. This is because the CE cells in the healthy rabbit cornea are already at a density of ∼3,000 cells/mm^2^, and thus it is encouraging that at the site of engraftment µMonolayers were able transiently increase this density to ∼5,000 cells/mm^2^ before migrating out over time (**Fig 7 and fig. S7**). Ideally, this technology would be tested in an in vivo model that is more representative of patients with a low CE cell density of <500 cell/mm^2^, our target population. However, such an in vivo model does not currently exist (*39*), as the low density observed in humans is related to genetic deposition, decades of aging, and exposure to UV light (*40, 41*). Developing an animal model to effectively recapitulate the human disease state we are ultimately aiming to treat is thus challenging. For these reasons, we chose the healthy rabbit in vivo model, as we determined it was more relevant than a complete injury and removal model that has been used in other rabbit CE studies. A limitation of this study that it is important to consider is the variability in cell density from cornea to cornea, even for individual rabbit. This adds a layer of difficulty to quantifying any effect on cell density within the healthy CE model. However, the results from our in vitro studies utilizing lower density CE monolayers showed a significant increase in cell density of >50% when μMonolayers were injected, as compared to single cell suspension controls (**Fig. 4**). This result, combined with the in vivo rabbit studies (**Fig. 6 and 7**), demonstrates shrink-wrapped CE cell μMonolayers can engraft into endothelial tissues at a high level that cannot be matched by any other approach.

As we take the next steps towards clinical translation, we will expand upon these proof-of-concept studies to include larger in vivo rabbit studies in both healthy and fully stripped endothelium models. While these models are not fully representative of patients with low CE cell density, the fully stripped endothelium model is in line with how CE cell replacement using single cells in suspension is done in current human clinical trials and will allow us to compare this technology more directly with those methods. In the future, we also hope to apply this technology to therapeutic cell delivery in other organs that suffer from donor shortages, such as the heart and liver where we could restore function through cardiomyocyte and hepatocyte delivery, respectively. There are also diseases where the unique ability of shrink-wrapped µMonolayers to engraft into epithelial layers could provide new therapeutic options, such as the delivery of airway cells genetically edited to restore cystic fibrosis transmembrane conductance regulator (CFTR) function for cystic fibrosis patients. Or for other forms of lung disease such as post COVID-19 infection, delivering alveolar type II epithelial cells to damaged distal regions. While there are many paths towards clinical impact, the key take home of this work is that it is possible to use our shrink-wrapping approach to engineer small clusters of cells that retain key phenotypic markers and are primed for rapid and enhanced engraftment into tissues, compared to existing alternatives.

## Materials and Methods

### Study Design

The research objectives of this study were to: 1) modify our previously published SHELL technique that was used to shrink-wrap single cells in order to shrink-wrap small islands of corneal endothelial cells into patches of endothelium termed µMonolayers, 2) to investigate whether shrink-wrapped μMonolayers would integrate into and increase cell density *in vitro* and 3) determine if the μMonolayers were capable of engrafting into an existing healthy CE *in vivo*. Primary bovine and rabbit cells were used throughout the study using previously published cell culture methods with minor modifications in the case of the rabbit cells (*30, 33*). Sample sizes for in vitro studies were determined by using the minimum number of samples to be considered statistically significant and time points/end points were based on our previously published studies. For the in vitro cell density study, 4 replicates per sample type per time point were used and one full study was completed. Data from day 3 time point was used to determine if the sample size was sufficient enough to provide statistical significance. At day 7 one control sample and at day 14 one single cells sample was lost during fixing and staining and therefore the n=3 for those sample types. In vivo studies were designed to be pilot studies and as such the number of rabbits per study was kept to 2-3 animals per time point and condition to establish repeatability.

### ECM scaffold fabrication

The ECM scaffolds were fabricated via previously described surface-initiated assembly techniques with minor modifications (*24, 25*). Briefly, 1 cm × 1 cm PDMS stamps designed to have 200 μm square features were fabricated via standard soft lithography techniques. The stamps were sonicated in 50% ethanol for 60 minutes, dried under a stream of nitrogen and incubated for 60 minutes with a 50:50 mixture of 50 μg/mL collagen IV (COL4) and 50 μg/mL laminin (LAM) (**Fig. 1 step 1**). Either 50% AlexaFluor 488 labeled COL4 or 50% AlexaFluor 633 labeled LAM (for a final concentration of 25% labeled protein) was used to visualize the pattern transfer. Following incubation, the stamps were rinsed in sterile water, dried under a stream of nitrogen, and brought into conformal contact with poly(N-isopropylacrylamide) (PIPAAm) (2% high molecular weight, Scientific Polymers) coated 18 or 25 mm glass coverslips for 30 minutes to ensure transfer of the squares (**Fig. 1 step 2**). ECM squares microcontact printed on PDMS coverslips were used as controls. Upon stamp removal, laser scanning confocal microscopy was used to determine the quality of the transferred ECM squares (Nikon AZ100).

### Corneal endothelial cell culture

Bovine CE cells were isolated and cultured as previously described (*30, 42*). Briefly, corneas were excised from the whole globe (Pel Freez), incubated endothelial side up in a ceramic 12 well spot plate with 400 μL of TrypLE Express for 20 minutes. The cells were then gently scraped from the cornea using a rubber spatula, centrifuged at 1500 RPM for 5 minutes, resuspended in 5 mL of culture media (low glucose DMEM with 10% FBS, 1% Pen/Strep/AmphB and 0.5% gentamicin, designated at P0 and cultured in a 50 kPa PDMS coated T-25 flask that was pre-coated with COL4 (*30*). Fifty whole eyes were received at a time and were used to seed 5 T-25 flasks. Cells were cultured until confluence and split 1:3 until they were used once confluent at P2.

Whole rabbit eyes were received on ice from Pel Freez Biologicals. Corneas were excised from the whole globe, the CE and Descemet’s Membrane were manually stripped with forceps and incubated in Dispase (1U/mL, Stem Cell Technologies) for 1.5 hours at 37°C to detach the rabbit CE cells (RCECs) from the Descemet’s Membrane. The RCECs were then gently pipetted up and down, diluted in culture media (DMEM/F12, 10% FBS, 0.5% Pen/strep), centrifuged at 1500 RPM for 5 minutes, resuspended in 10 mL of culture media, designated at P0 and cultured COL4 coated T-25 flasks with the equivalent of 15-25 eyes per flask depending on cell yield. RCECs were cultured until confluence and split 1:2 and used in all experiments once confluent at P1 or P2.

### Shrink-wrapping CE Cell µMonolayers in ECM scaffolds

Patterned coverslips were secured with vacuum grease to the bottom of 35 mm petri dishes which were placed on a dry block set to 52°C. This resulted in the coverslips reaching (within 30 min) and holding at 40°C. Bovine CE cells were released from the culture flask with TrypLE Express, centrifuged and resuspended at a density of 150,000 cells/mL in 15 mL centrifuge tubes. The tubes were placed in a dry block set at 45 °C for approximately 5 minutes, or until the cell solution just reached 40 °C and 2 mL of cell suspension was added to each 35 mm dish before it was immediately placed in an incubator (37 °C, 5% CO2). Cells were cultured for 24 hours to allow them to form μMonolayers on the 200 μm squares. Samples were then removed from the incubator, rinsed twice in 37 °C media to remove non-adherent single cells, 2 mL of fresh warm media was added, and the sample was allowed to cool to room temperature. Once the temperature decreased to less than 32 °C the PIPAAm dissolved and released the µMonolayers. The release process was recorded using a Photometrics CoolSnap camera. Following release, the µMonolayers were collected via centrifugation at 1500 rpm for 5 minutes before use in further experiments. CE cells seeded on to PDMS coverslips were used as a control.

### Immunostaining of shrink-wrapped CE Cell μMonolayers

Shrink-wrapped µMonolayers resuspended in PBS containing Ca^2+^ and Mg^2+^ (PBS++) were injected through a small gauge needle onto a glass coverslip and allowed to settle for ∼15 minutes before fixation for 15 minutes in 4% paraformaldehyde in PBS++. Samples were gently washed 2 times with PBS++ and incubated with 1:100 dilution of DAPI, 1:100 dilution of mouse anti-ZO-1 antibody (Life Technologies) and 3:200 dilution of AlexaFluor 488. Samples were rinsed 2 times for 5 minutes with PBS++ and incubated with 1:100 dilution of AlexaFluor 555 goat anti-mouse secondary antibody for 2 hours. Samples were rinsed 2 times for 5 minutes with PBS++, mounted on glass slides with Pro-Long Gold Antifade (Life Technologies) and then imaged on a Zeiss LSM 700 confocal microscope.

### Viability of shrink-wrapped CE cells μMonolayers post-injection

After centrifugation, shrink-wrapped µMonolayers or TrypLE Express released single cells were resuspended in 200 µL of growth media, drawn up into a 28G needle, injected into a petri dish and incubated with 2 µM calcein AM and 4 µM EthD-1 (Live/Dead Viability/Cytotoxicity Kit, Life Technologies) in PBS++ for 30 minutes at 37 ° C. After 30 minutes, samples were imaged on a Zeiss LSM 700 confocal; 5 images per sample and 3 samples per type were used. The number of live and dead cells was counted manually using ImageJ’s multi-point tool. The number of live cells was divided by the number of total cells to determine the percent viability of both the shrink-wrapped cells and enzymatically released cells. The data was compared using a Student’s t-test in SigmaPlot. The same methods were used to test the viability of the cells through a 34G needle to test the smallest needle that could be used.

### Seeding of shrink-wrapped CE Cell μMonolayers and single CE cells on stromal mimics

Self-compressed collagen type I films were prepared as previously described to mimic the structure of the underlying stroma (*42*). Briefly, a 6 mg/mL collagen type I gel solution was prepared per manufacturer’s instructions and pipetted into 9 mm diameter silicone ring molds on top of glass coverslips. The gels were placed into a humid incubator (37 °C, 5% CO2) for 3 hours to compress under their own weight. The gels were then dried completely in a biohood followed by rehydration in PBS, forming a thin collagen type I stromal mimic. Shrink-wrapped CE cell µMonolayers were seeded onto the films at a 1:1 ratio of stamped coverslip to collagen type I film. As a control, CE cells that were cultured in the flasks and enzymatically released using TrypLE Express into a single cell suspension were seeded onto collagen type I films. The number of control single cells seeded was equal to the number of cells seeded in the μMonolayers, assuming each of the 200 μm ECM squares used for shrink-wrapping was completely covered in cells. The average number of cells on the 200 μm ECM squares was 30 cells, so with 1600 squares per stamp we seeded ∼48,000 cells per sample. Therefore, 50,000 cells per sample were seeded for the controls. At 6 and 24 hours, samples were removed from culture and fixed and stained for the nucleus, ZO-1 (tight junction protein) and F-actin. Briefly, samples were rinsed 2 times in PBS++, fixed in 4% paraformaldehyde in PBS++ with 0.05% Triton-X 100 for 15 minutes. Samples were rinsed 2 times for 5 minutes with PBS++ and incubated with 5 drops of NucBlue (Life Technologies) for 10 minutes. Samples were rinsed once with PBS++ and incubated with 1:100 dilution of mouse anti-ZO-1 antibody (Life Technologies) and 3:200 dilution of AlexaFluor 488 or 633 phalloidin for 2 hours. Samples were rinsed 3 times for 5 minutes with PBS++ and incubated with 1:100 dilution of AlexaFluor 555 goat anti-mouse secondary antibody for 2 hours. Samples were rinsed 3 times for 5 minutes with PBS++, mounted on glass slides using Pro-Long Gold Anti-fade and imaged on a Zeiss LSM 700 confocal microscope.

### In vitro integration of shrink-wrapped μMonolayers vs single bovine CE cells

To mimic a low-density CE, 25,000 P5 bovine cells were seeded onto the collagen type I stromal mimics, as described above, until confluent to form the low-density monolayers. Shrink-wrapped µMonolayers and single CE cells were prepared as above, labeled with CellTracker Green (Life Technologies) for 30 minutes, centrifuged, diluted to the equivalent of 50,000 cells/sample and injected onto the low-density monolayers. Low density monolayers with no cells injected on top served as controls. Samples were rinsed 3 hours post injection to mimic the in vivo procedures and new media was added. Media was changed every two days thereafter. Samples were fixed and stained at days 3, 7 and 14 as described above. A Zeiss LSM700 confocal was used to image 10 random spots on each sample and the cell density was manually counted using the multi-point selection tool in ImageJ to count cell nuclei. The number of nuclei was divided by the image area to obtain the cells/mm^2^ per image. The cell density for each sample was determined by averaging the cell densities of each image and the average cell density of each sample type was determined by averaging the cell density of the 3-4 samples. The data was compared using a one-way ANOVA on ranks with Tukey’s test (day 3) or one-way ANOVA (days 7 and 14) with Tukey’s Test is SigmaPlot. To examine the outgrowth of the shrink-wrapped µMonolayers over time, confocal images centered around and individual shrink-wrapped µMonolayer (day 3 n=33, day 7 n=37, day 14 n=40) were collected and the CellTracker channel was converted into a binary black and white image. The binary images for each sample type were then converted into one Z-stack and analyzed via the Heat Map for Z-stacks plugin (relative without log10) for ImageJ to determine the average pixel density of CellTracker.

### Live imaging of in vitro integration of shrink-wrapped bovine CE cells

For live imaging, the monolayer on the collagen type I stromal mimic was first incubated for 30 minutes with CellTracker Orange to differentiate between the existing monolayer and injected cells, which were labeled with CellTracker Green as described above. HEPES buffered Opti-MEM I Reduced Serum Media (Life Technologies) with 10% FBS, 1% Pen/Strep was added to the monolayer and shrink-wrapped µMonolayers that were prepared as described above were injected through a 30G needle on top of the sample. The sample was placed on the Zeiss LSM700 confocal equipped with a temperature chamber set to 37°C for 30 minutes to allow for the cells to settle. Using the Definite Focus system, a time-lapse series of one z-stack was obtained every hour for 48 hours. Videos from the time-lapse images were created using the Imaris Software.

### Ex vivo integration of shrink-wrapped rabbit CE cells

Rabbit eyes were placed cornea up in a 12-well plate and shrink-wrapped RCEC µMonolayers were prepared as described above. Two samples of µMonolayers per ex vivo eye were prepared and resuspended in 100 µL of DMEM/F12. A 30-G insulin syringe was used to draw up the full 100 µL, the needle was inserted into the center of the cornea until it was visible in the anterior chamber and 5 0µL of the suspension was injected. This resulted in the equivalent of 50,000 cells injected into the anterior chamber. The needle was held in place for a few seconds to ensure the media and cells did not come back out of the injection site. The injection was viewed under a stereomicroscope and the pink color of the media filling the anterior chamber was visible, indicating successful injection. The eyes were flipped and incubated cornea down for 3 hours at 37°C, 5% CO_2_ in a humidified incubator. After 3 hours, the whole eye was placed in 2% paraformaldehyde (PBS++) at 4 °C for 24 hours. After 24 hours the eye was rinsed in PBS and the cornea was excised and rinsed 3 times for 5 mins. The cornea was then incubated CE facing down on 1 mL of PBS++ containing 2 drops of NucBlue (Life Technologies), 2:100 dilution of mouse anti-ZO-1 antibody (Life Technologies) and 3:200 dilution of AlexaFluor 488 Phalloidin (Life Technologies) for 2 hours at room temperature. Corneas were then rinsed 3 times for 5 minutes in PBS followed by a 2-hour incubation on 1mL PBS++ with 2:100 dilution of AlexaFluor 555 goat anti-mouse secondary antibody for 2 hours and stored in PBS before imaging on the Zeiss LSM700 confocal.

### In vivo injection and integration of shrink-wrapped CE cells

All experimental procedures were reviewed and approved by the University of Pittsburgh Institutional Animal Care and Use Committee (IACUC) and carried out according to guidelines of the Association for Research in Vision and Ophthalmology Resolution on the Use of Animals in Ophthalmic and Vision Research. For both in vivo experiments, shrink-wrapped RCEC µMonolayers were prepared as described above with one minor modification: cells were labeled with Vybrant DiO 1 day prior to seeding on to the ECM nano-scaffolds by incubating cells in 1mL of media with 5 µL of Vybrant DiO for 30 minutes followed by three, 10-minute rinses with fresh media. An excess number of µMonolayer samples were prepared to ensure there was enough volume for injection. The shrink-wrapped µMonolayers were released as described above and after centrifugation at 1500 rpm for 5 min, the shrink-wrapped µMonolayers were resuspended in DMEM/F12 at the equivalent of 100,000 cells per 50 μL injection volume (2 stamped samples per 50 μL).

For the first experiment, control single cells were prepared as described above and resuspended in DMEM/F12 at a density of 100,000 cells in a 50 μL injection volume. Six female New Zealand white rabbits with healthy intact CEs weighing approximately 2.5kg were used for this study. Rabbits were anesthetized with Ketamine (40 mg/kg) and Xylazine (4 mg/kg) intramuscular injection followed by isoflurane inhalation to keep rabbits under sedation for 3 hours. One rabbit did not survive the anesthetization. Rabbits #1 & 2 were injected in the right eye with 50 µL (∼100,000 cells) of the single cell suspension. Rabbits #3, 4 and 5 were injected with 50 µL of the shrink-wrapped μMonolayer suspension into the right eye using a 30G needle attached to a 500 µL syringe. A tunnel in the corneal stroma was made for the injecting which prevented cell leakage after injection. Immediately after injection, each rabbit was placed on their side with the injected eye facing down for 3 hours to ensure attachment of the cells. On day 7, rabbits were anesthetized with an intramuscular injection of ketamine (40 mg/kg) and xylazine (4 mg/kg) and then euthanized with of Euthasol solution (1 mg per 4 lbs) containing (390 mg/mL Sodium Pentobarbitol, 50 mg/mL Phenytoin Sodium) through an ear vein injection. Photographic images were obtained via the Google Pixel 2 camera (fig. S3) to document eye clarity and the endothelium was viewed using a Nidek Confoscan 3 (fig. S4 Eyes were then immediately enucleated and intravitreally injected with 100 µL 2% paraformaldehyde in PBS++. The whole eye was then immersed in 2% paraformaldehyde in PBS++ and fixed at 4°C for 24 hours. After 24 hours the eye was rinsed in PBS and the cornea was excised and rinsed 3x’s for 5 mins. The cornea was then incubated CE facing down on 1mL of PBS++ containing 2 drops of NucBlue (Life Technologies) and 2:100 dilution of mouse anti-ZO-1 antibody (Life Technologies) for 2 hours at room temperature. Corneas were then rinsed 3 times for 5 minutes in PBS followed by 2-hour incubation on 1mL PBS++ with 2:100 dilution of AlexaFluor 555 goat anti-mouse secondary antibody for 2 hours and stored in PBS before imaging on the Zeiss LSM700 confocal or a Nikon FN1 base with an A1R HD MP Confocal module. To quantify the density of the integrated cells, the green cells were manually traced with the freehand selection tool in ImageJ, and the “Measure” function was used to determine the area. The multi-point selection tool was used to determine the number of nuclei within that area and the density was determined by dividing the number of nuclei by the area. The rectangle tool was used to select control areas, areas of non-green (i.e. non-DiO labeled) cells, elsewhere in the image of similar area. The area and cell number were determined the same way as for the DiO labeled areas and the cell density was calculated by dividing the number of nuclei by the area. The number of control areas per image was matched to the number of DiO labeled areas (fig. S5). Data from the injected eye of both rabbits was pooled together and the cell density of the areas with green cells were statistically compared to those without SigmaPlot using a student t-test.

For the second experiment, only shrink-wrapped μMonolayers were used to determine if cells would integrate and remain in the CE long-term. Six female New Zealand white rabbits with healthy intact CEs weighing approximately 2.5kg were used for this study. Rabbits were anesthetized as described above. All rabbits were anesthetized and injected with 50 µL of the shrink-wrapped μMonolayer suspension into the right eye. Immediately after injection, each rabbit was placed on their side with the injected eye facing down for 3 hours to ensure attachment of the cells. One rabbit went into tachycardia right at the end of the 3 hours and was revived, however, it suffered brain damage as a result of the amount of time without oxygen. That rabbit was sacrificed 24 hours post-injection and the eye was removed and processed as described above and used to determine if the injection processed had been successful (data not shown). At 14 days (2 rabbits) or 28 days (3 rabbits) post-injection, rabbits were sacrificed (as described above) and photographic images were obtained via the Google Pixel 2 camera (fig. S3) to document eye clarity and the endothelium was viewed using a Nidek Confoscan 3 (fig. S4 Following imaging, eyes were immediately enucleated and intravitreally injected with 100 uL 2% paraformaldehyde in PBS++. The whole eye was then immersed in 2 % paraformaldehyde in PBS++ and fixed at 4 °C for 24 hours. After 24 hours the eye was rinsed in PBS and the cornea was excised and rinsed 3x’s for 5 mins. The cornea was then incubated CE facing down on 1mL of PBS++ containing 2 drops of NucBlue (Life Technologies) and 2:100 dilution of mouse anti-ZO-1 antibody (Life Technologies) for 2 hours at room temperature. Corneas were then rinsed 3 times for 5 minutes in PBS followed by 2-hour incubation on 1mL PBS++ with 2:100 dilution of AlexaFluor 555 goat anti-mouse secondary antibody for 2 hours and stored in PBS before imaging on the Zeiss LSM700 confocal or a Nikon FN1 base with an A1R HD MP Confocal module. To quantify the cell density, images with DiO labeled cells present were taken (n=10+ images) and images were taken far away in areas where there were no green cells (n=5 images) in each cornea. Because the DiO labeled may have faded over time, in this case the cell density of the entire image was counted using the number of nuclei (counted manually via the multi-point tool with only full nuclei being counted) divided by the area of the image. To statistically compare the data, for each rabbit, the density of the images with green cells was compared to the density of the images with no green cells in SigmaPlot using student T-test.

## Supplementary Materials

Movie S1. Release of Shrink-wrapped μMonolayers

Movie S2. Time-lapse In Vitro Integration of Shrink-wrapped μMonolayers (Top-down view)

Movie S3. Time-lapse In Vitro Integration of Shrink-wrapped μMonolayers (Side view)

Figure S1. AFM showing nanostructure and height of the patterned ECM nanoscaffolds.

Figure S2. Schematic showing the ex vivo experimental setup.

Figure S3. Rabbit eye photographs.

Figure S4. Confoscan showing normal cobblestone morphology of corneas post injection at 1 and 4 weeks.

Figure S5. Examples of equivalent areas used for Native Cell Density calculations.

Figure S6. Large area Tile-scans of In Vivo Corneas 1 week Post-Injection.

Figure S7. Cell density of areas with green DiO labeled cells compared to areas with no labeled cells

## Author contributions

RP completed all in vitro and ex vivo experiments, prepped all of the cells for and observed the in vivo studies, completed all data analysis, prepared all the figures, and prepared the manuscript. YD performed the injections for the in vivo experiments, gave feedback on data and figures, and editing of the manuscript. MG did all rabbit anesthetization and oversaw all in vivo rabbit experiments and care as well as did all rabbit experiment post-prep and enucleation. SC assisted with rabbit eye dissection for cell retrieval. DS acquired the multi-photon images of the rabbit corneas for in vivo experiments. IK assisted with rabbit experiments. JF and AF oversaw the entire study, gave feedback on data and figures and editing/preparation of the manuscript.

## Data and materials availability

All raw data is available upon written request to the authors. Anyone requesting materials related to shrink-wrapping cells must complete an MTA in order receive those materials.

## Movie Captions

**Movie S1. Release of Shrink-wrapped μMonolayers**. A time-lapse video showing the release of the shrink-wrapped μMonolayers. Warm PBS with calcium and magnesium was added to the sample and as the temperature decreases below the LCST of PIPAAm, it dissolves, resulting in the release and shrink-wrapping of the μMonolayers.

**Movie S2. Time-lapse In Vitro Integration of Shrink-wrapped μMonolayers (Top-down view)**. This is a time-lapse confocal microscopy video from the top-down view showing the integration of the shrink-wrapped μMonolayers (labeled with CellTracker Green) into an existing monolayer of CE cells (labeled with CellTracker Orange, appearing RED). The micropatterned ECM used to shrink-wrap the μMonolayers is shown in purple.

**Movie S3. Time-lapse In Vitro Integration of Shrink-wrapped μMonolayers (Side view)**. This is a time-lapse confocal microscopy video rendered from the side view showing the integration of the shrink-wrapped μMonolayers (labeled with CellTracker Green) into an existing monolayer of CE cells (labeled with CellTracker Orange, appearing RED). The micropatterned ECM used to shrink-wrap the μMonolayers is shown in purple.

