## Supplemental Figures for "Injury-Free In Vivo Delivery and Engraftment into the Cornea Endothelium Using Extracellular Matrix Shrink-Wrapped Cells"

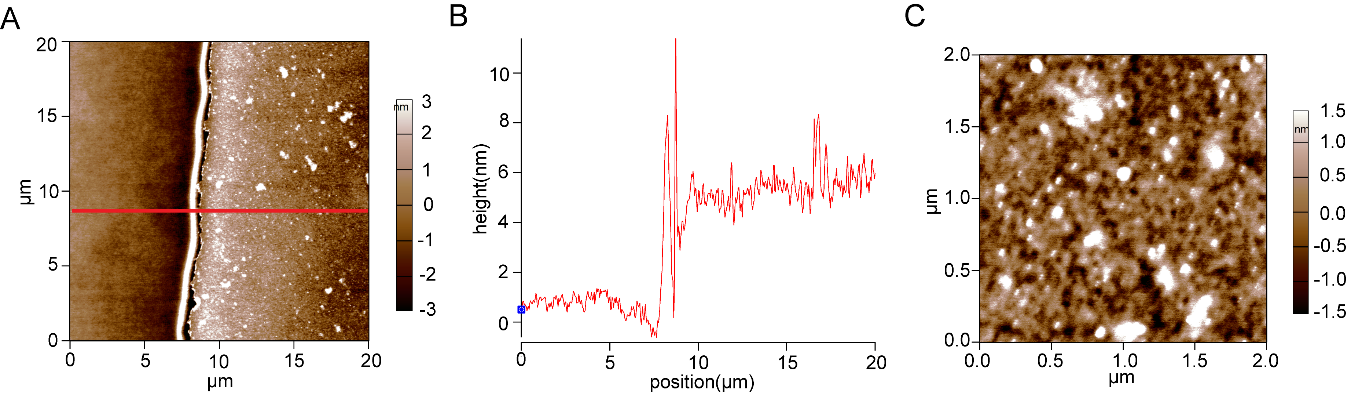

**Supplemental Figure S1.** **AFM images showing nanostructure and height of the patterned ECM nanoscaffolds.** (A) AFM scan of the side of an ECM nanoscaffold. (B) Height trace of the red line shown in A showing that at the very edge the scaffold is approximately 10 nm high, but that the scaffold is between 4-6 nm across the rest of the scaffold. (C) Zoomed in AFM scan of the patterned ECM showing the basement membrane like structure of the scaffold. Images were acquired on an Asylum Research MFP-3D-BIO AFM in AC in air mode using AC-160 cantilevers.

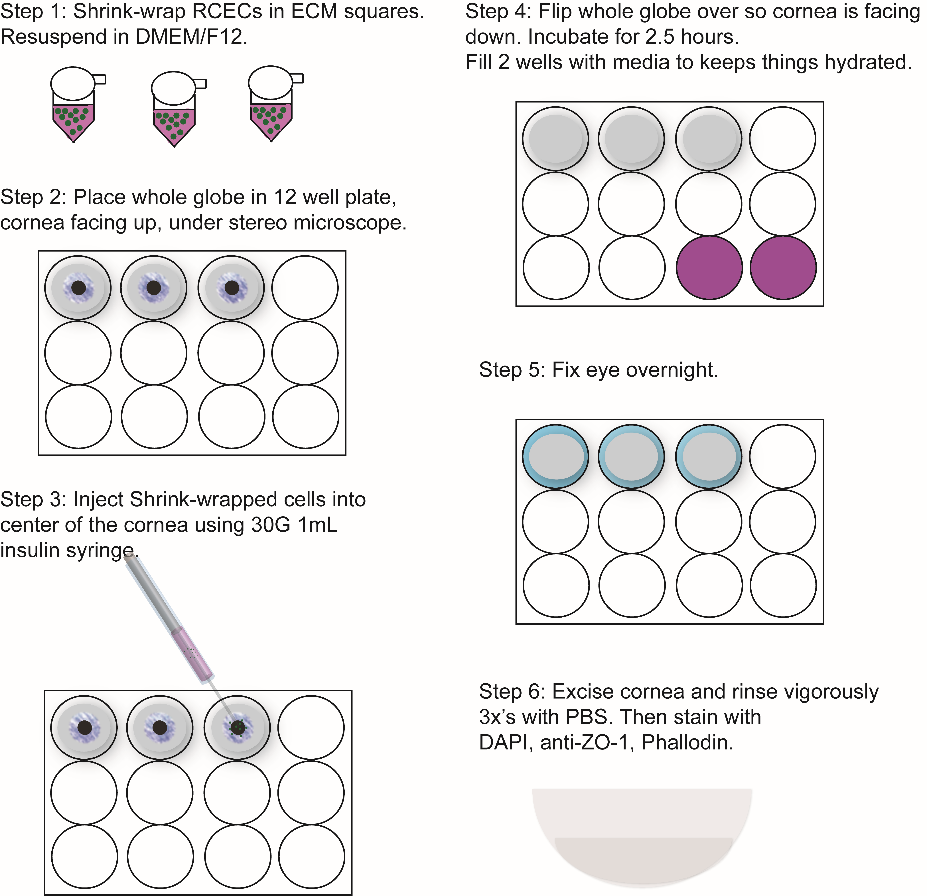

**Supplemental Figure S2:** **Schematic showing the ex vivo experimental setup.** In Step 1, shrink-wrapped rabbit CE cells (RCECs) were suspended in sterile DMEM/F12 media. In Step 2, Whole enucleated rabbit eyes (i.e., globes) were placed cornea side up in a 12 well plate. In Step 3, the shrink-wrapped RCECs from Step 1 were injected into the anterior chamber to simulate the clinical process of cell injection for CE repair. In Step 4, the globes were flipped upside down with the cornea facing the bottom of the well. This was done so that the injected shrink-wrapped cells would contact the posterior surface of the cornea through gravitational settlement. In Step 5, the globes were fixed in paraformaldehyde overnight. In Step 6, the corneas were cut out from the globes, rinsed thoroughly, and then immunofluorescently stained for nuclei, ZO-1, and F-actin.

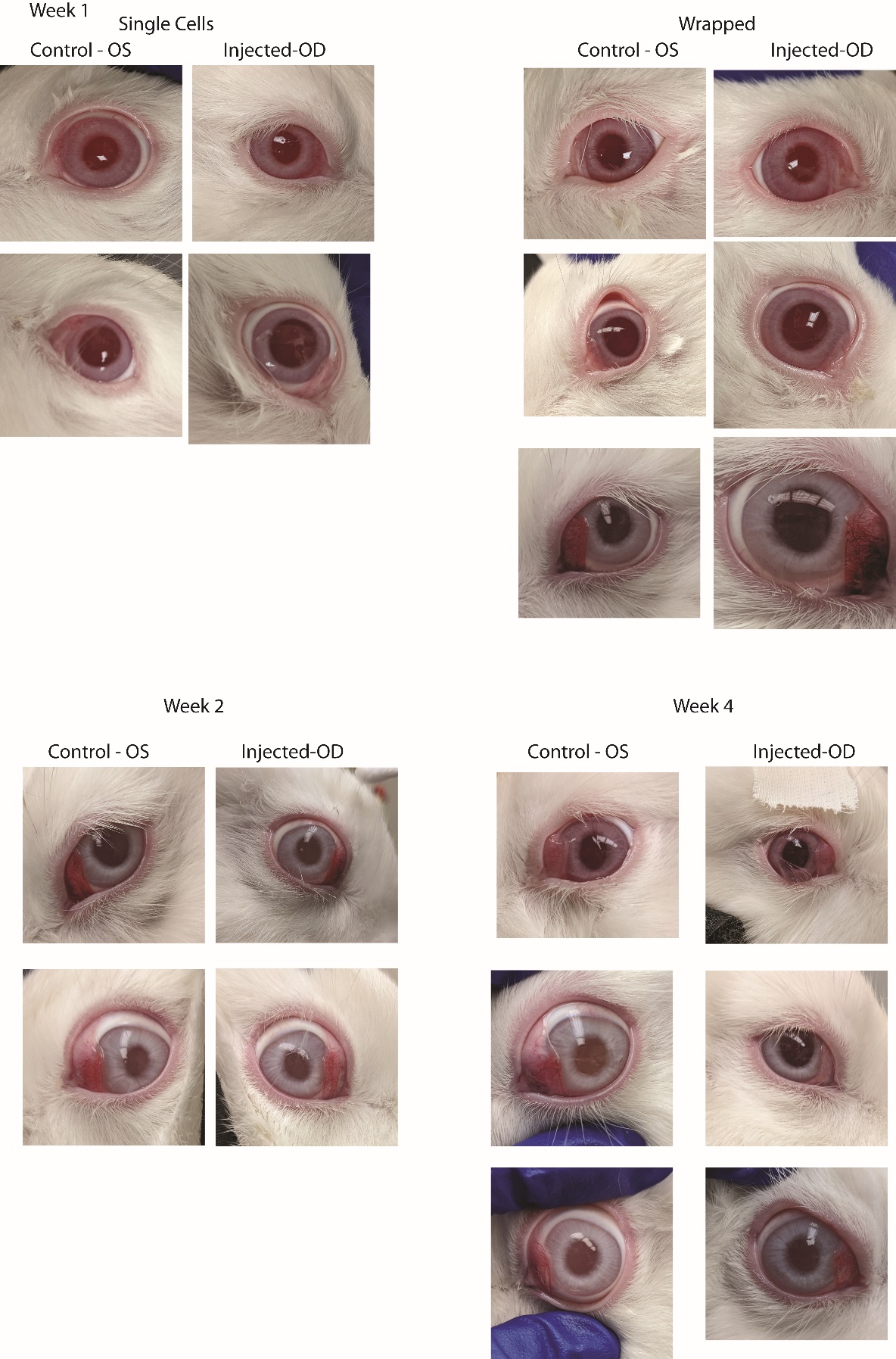

**Supplemental Figure S3:** **Rabbit eye photographs.** Images showing the uninjected control eye (OS) of each rabbit, compared to the injected (OD) eye of each rabbit. All eyes were clear at all time points showing no cloudiness or obvious irritation from the injection of single or shrink-wrapped μMonolayer cells.

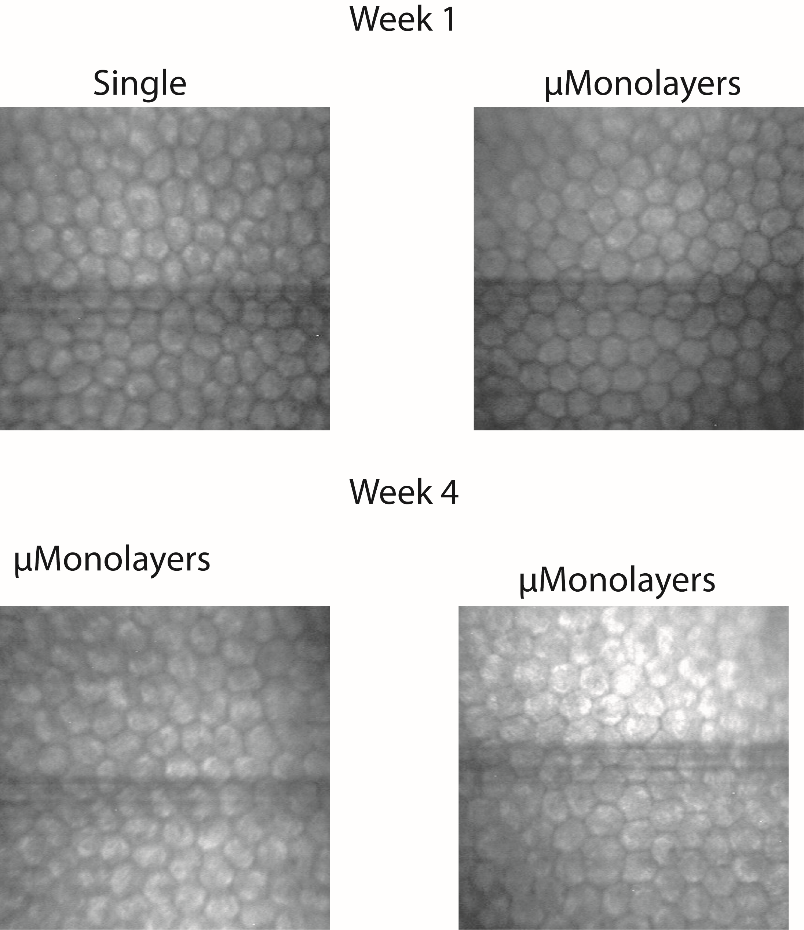

**Supplemental Figure S4:** **Confoscan showing normal cobblestone morphology of corneas post injection at 1 and 4 weeks.** Rabbit eyes injected with DiO labeled endothelial cells, either single cells or shrink-wrapped as µMonolayers, were imaged with the Confoscan at 1 and 4 weeks. The endothelium appeared normal, with the expected cobblestone morphology and cell density.

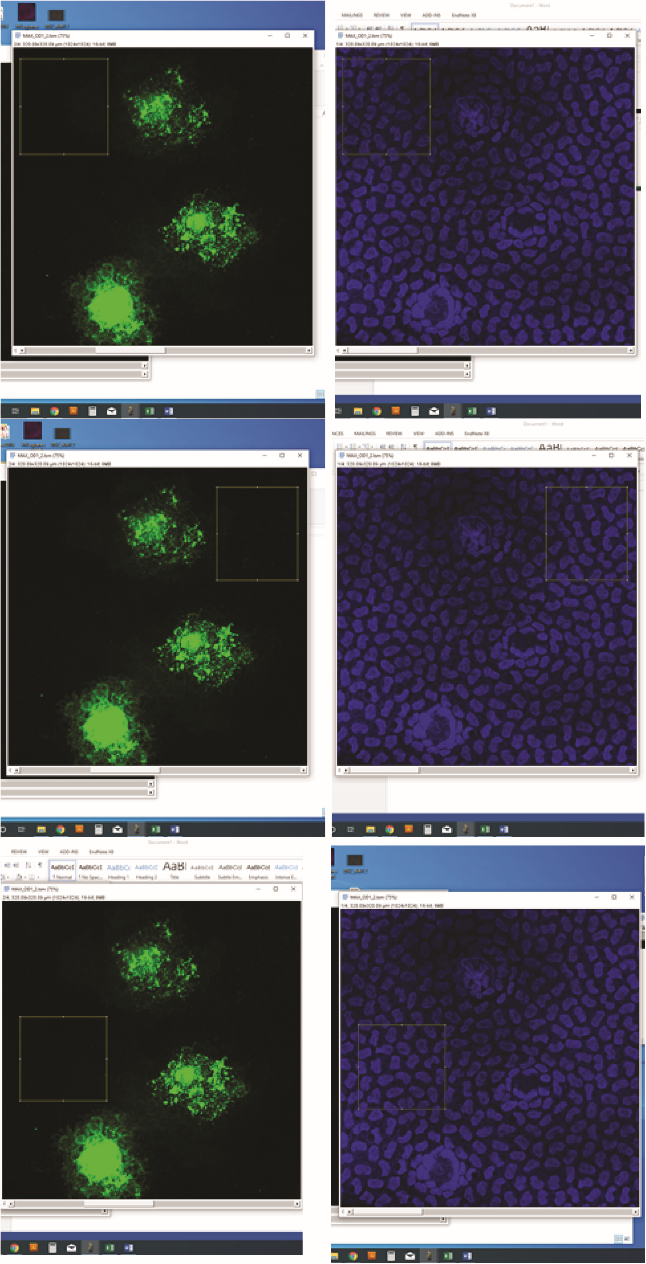

**Supplemental Figure S5:** **Examples of equivalent areas used for cell density calculations.** For the DiO labled areas, the cells were traced with the pen feature in ImageJ, the nuclei were manually counted and the area was determined by ImageJ. To get the native density the areas shown above in the boxes in yellow were used. The same number of equivalent areas as DiO labeled areas were counted per image and the areas were designed to be approximately the same size as the DiO labeled regions.

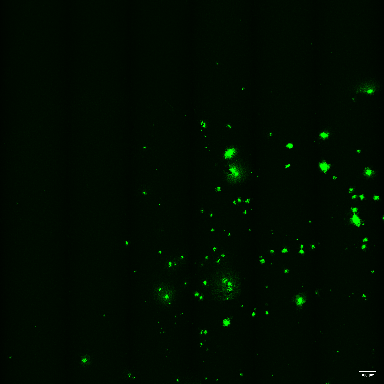

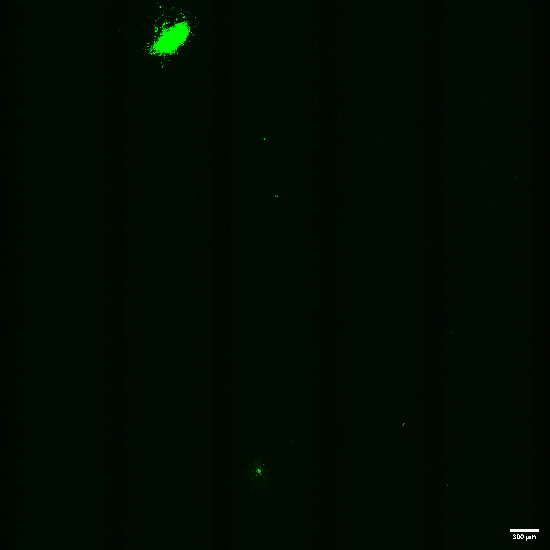
**A**	**B**

**Supplemental Figure S6: Large area Tile-scans of In Vivo Corneas 1 week Post-Injection.** (A) A representative image of the cornea with single cells injected (5 X 5 mm tile scan). The large spot is the area of the injection and some green signal is due to light reflection from the damage to the stroma. (B) A representative image of DiO positive μMonolayers present in the corneal endothelium 1 week post-injection (5 X 5 mm tile scan). Scale bars = 300 μm

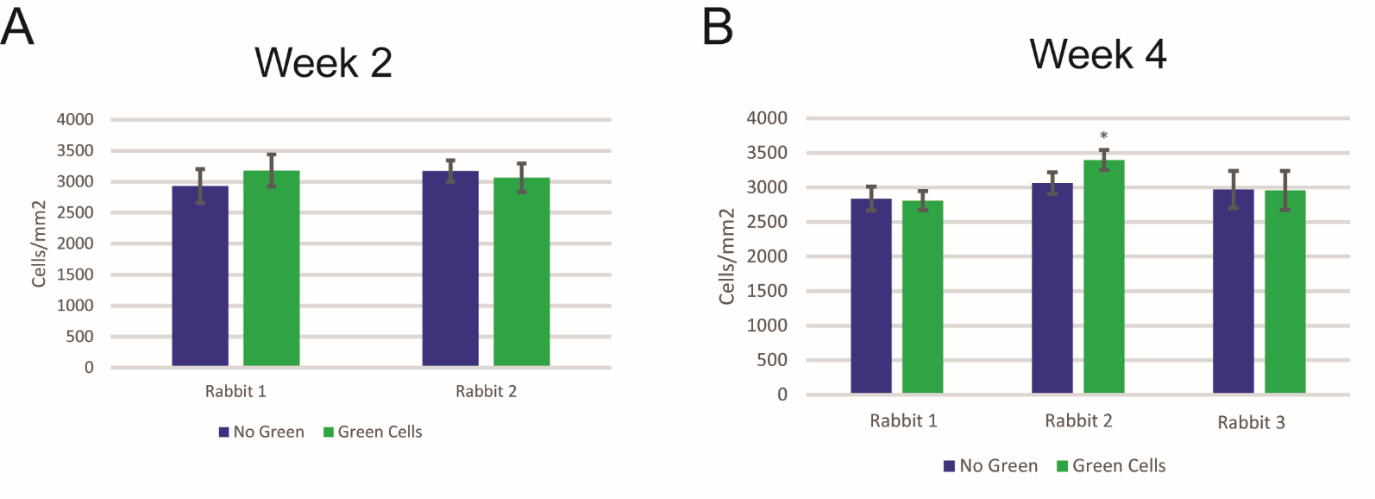

**Supplemental Figure 7:** **Cell density of areas with green DiO labeled cells compared to areas with no labeled cells.** To quantify the cell density of the rabbit CEs post injection, images in areas with DiO labeled cells present were taken (n=10+ images) and images were taken far away in areas where there were no green cells (n=5 images) in each injected cornea. Because the DiO labeled may have faded over time, in this case the cell density of the entire image was counted using the number of nuclei (counted manually via the multi-point tool with only full nuclei being counted) divided by the area of the image. To statistically compare the data, for each rabbit cornea, the average density of the images with green cells was compared to the average density of the images with no green cells in SigmaPlot using student T-test. Data is represented as mean ± standard deviation and * indicates statistically significant p<0.05.
